# A Multidimensional Array Representation of State-Transition Model Dynamics

**DOI:** 10.1101/670612

**Authors:** Eline M. Krijkamp, Fernando Alarid-Escudero, Eva A. Enns, Petros Pechlivanoglou, M.G. Myriam Hunink, Hawre J. Jalal

**Author notes:** Contributed equally to this work (Eline M. Krijkamp, Fernando Alarid-Escudero).

## Abstract

Cost-effectiveness analyses often rely on cohort state-transition models (cSTMs). The cohort trace is the primary outcome of cSTMs, which captures the proportion of the cohort in each health state over time (state occupancy). However, the cohort trace is an aggregated measure that does not capture the information about the specific transitions among the health states (transition dynamics). In practice, these transition dynamics are crucial in many applications, such as incorporating transition rewards or computing various epidemiological outcomes that could be used for model calibration and validation (e.g. disease incidence and lifetime risk). In this manuscript we propose modifying the transitional cSTMs cohort trace computation to compute and store cSTMs dynamics that capture both state occupancy and transition dynamics. This approach produces a multidimensional matrix from which both the state occupancy and the transition dynamics can be recovered. We highlight the advantages and potential applications of this approach with an example coded in R to facilitate the implementation of our method.

---

State transition models (STM) are decision models commonly used in cost-effectiveness analysis (CEA) to capture economic and health outcomes of different strategies over time in discrete time cycles [1, 2]. In a cohort STM (cSTM), the disease dynamics are captured by distributing a closed cohort among a mutually exclusive and collectively exhaustive set of health states [2, 3, 4]. The cohort trace is the primary outcome of cSTMs, which stores the proportion of the cohort in each health state over time (i.e., it summarizes state occupancy) [5, 1]. A limitation of the cohort trace is that it does not keep track of the transitions among health states over time (i.e., the transition dynamics of the cohort). As a consequence, it can only be used to capture outcomes that involve residing in a state for a full cycle to compute the so-called *state rewards* and does not contain a mechanism to assign transition rewards, which are applied only when specific transitions occur. It also limits the type of epidemiological outcomes that can be obtained from cSTMs. For example, obtaining incidence of a disease requires knowledge of the proportion of the population transitioning from a subset of states without disease to the state(s) representing the disease of interest.

To overcome the limitations of the cohort trace, we propose a multidimensional array-based approach that serves as a full summary of cSTM dynamics that complements the already useful cohort trace. The proposed approach allows modelers to efficiently calculate all measures of interest that rely on transition dynamics and at the same time to aggregate this into a standard cohort trace.

We start by providing a formal definition of cSTMs elements and the cohort trace. We complement this standard notation with a description of the detailed transition dynamics. Then, we introduce the multidimensional-array structure and show how it can be easily generated. Finally, we illustrate its use to compute a measure of interest that depends on transitions among health states. We demonstrate this approach with an illustrative example of a cSTM programmed in R provided in the supplementary material and in GitHub https://github.com/DARTH-git/state-transition-model-dynamics)[9, 10].

## Traditional cohort trace approach

We denote the distribution of the cohort across *n_s_* health states in a cSTM at cycle *t* as the state vector **m**_*t*_ of dimensions 1 × *n*_*s*_. That is, each element in **m**_*t*_ represents the proportion of the cohort in health state *i* at time *t*. Thus, **m**_*t*_ is written as

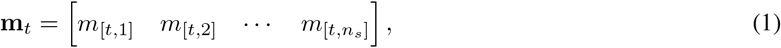

where the initial state vector **m**_0_ contains the distribution of the cohort across all *n*_*s*_ health states at the start of the simulation. The cohort distribution evolves over time governed by state transition probabilities. The probability of transitioning from health state *i* to health state *j* in cycle *t* is denoted as *p*_[*i,j,t*]_. The collection of transition probabilities across the model states over the time horizon forms the time-dependent state transition probability matrix, *P*_*t*_ of dimensions *n*_*s*_ × *n*_*s*_,

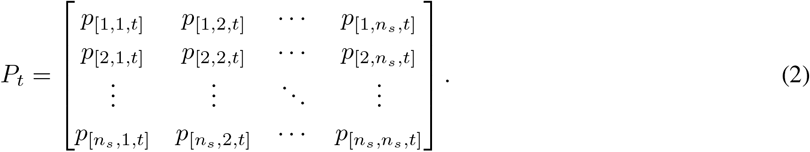

For any *t*, all rows of *P*_*t*_ must sum to one. Note that if *P*_*t*_ is equal for all *t* times, equation (2) becomes a time-homogeneous transition probability matrix, where *P*_*t*_ = *P*.

The state vector at cycle *t* + 1, **m**_*t*+1_, is then obtained by the inner product between the state vector at cycle *t*, **m**_*t*_, and the corresponding transition probability matrix *P*_*t*_, such that

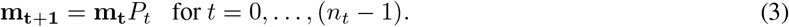

Stacking the state vectors by rows for all *t* = 0, …, *n*_*t*_ results in the full cohort trace matrix, *M*, of dimensions (*n*_*t*_ + 1) × *n*_*s*_, where each row is a state vector (–**m**_*t*_–), resulting in

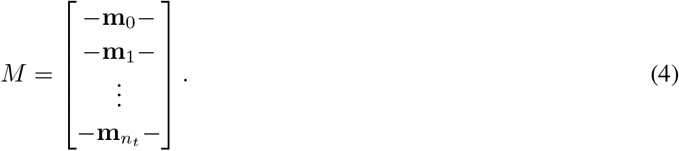

Together, the state vectors **m**_*t*_, the transition probability matrices *P*_*t*_ and the cohort trace *M* in equations (1), (2) and (4), respectively, represent the three main components of a cSTM.

## Dynamics-array approach

The trace matrix *M* aggregates transitions from all the states to a specific state, thus loses details of the transition dynamics. We propose to use a multidimensional array, **A**, of dimensions *n*_*s*_ × *n*_*s*_ × (*n*_*t*_ + 1) to store the proportion of the cohort that transitions between any two health states in each cycle over the time horizon. This array can be thought of as a set of two-dimensional matrices stacked along a third dimension that represents time. Below, we illustrate how to compute **A** from the three main components of cSTMs described above.

*A*_0_ represents the first “slice” of **A**. We compute *A*_0_ as a matrix containing the initial state vector **m**_0_ in its diagonal and 0s in the off-diagonal, such that

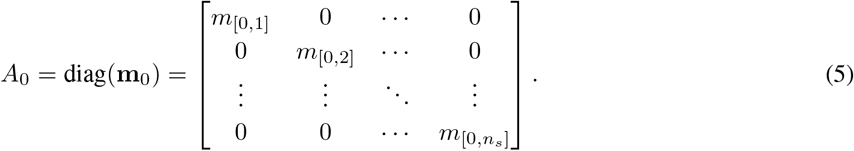

Each subsequent (*t* + 1)-th “slice” of **A** is obtained by multiplying a diagonal matrix of **m**_*t*_, denoted as diag(**m**_*t*_), by *P*_*t*_, such that

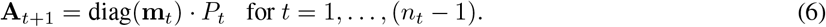

The resulting elements of the “*t*-th slice” of **A**, *A*_*t*_ for *t* > 0, are

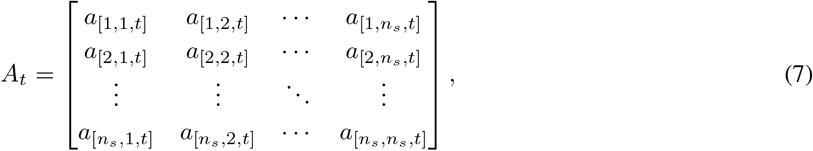

where *a*_[*i,j,t*]_ is the proportion of the cohort that transitions from state *i* to state *j* between cycles *t* − 1 and *t*, generated via

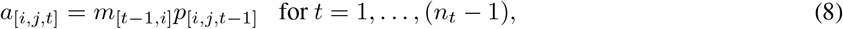

where *m*_[*t*−1,*i*]_ is the proportion of the cohort in state *i* at cycle *t* − 1 and *p*_[*i,j,t*−1]_ the corresponding transition probability at cycle *t* − 1 of transitioning from state *i* to state *j*. In other words, **A** stores the transition dynamics of a simulated cohort in a cSTM.

Figure 1 compares graphically the computation involved in both the traditional cohort-trace approach (a) and the dynamics-array approach (b) and shows the structures of the resulting cohort trace *M* and dynamics-array **A** (c) and how **A** recovers the transition dynamics that are being aggregated in *M*.

**Figure 1:**
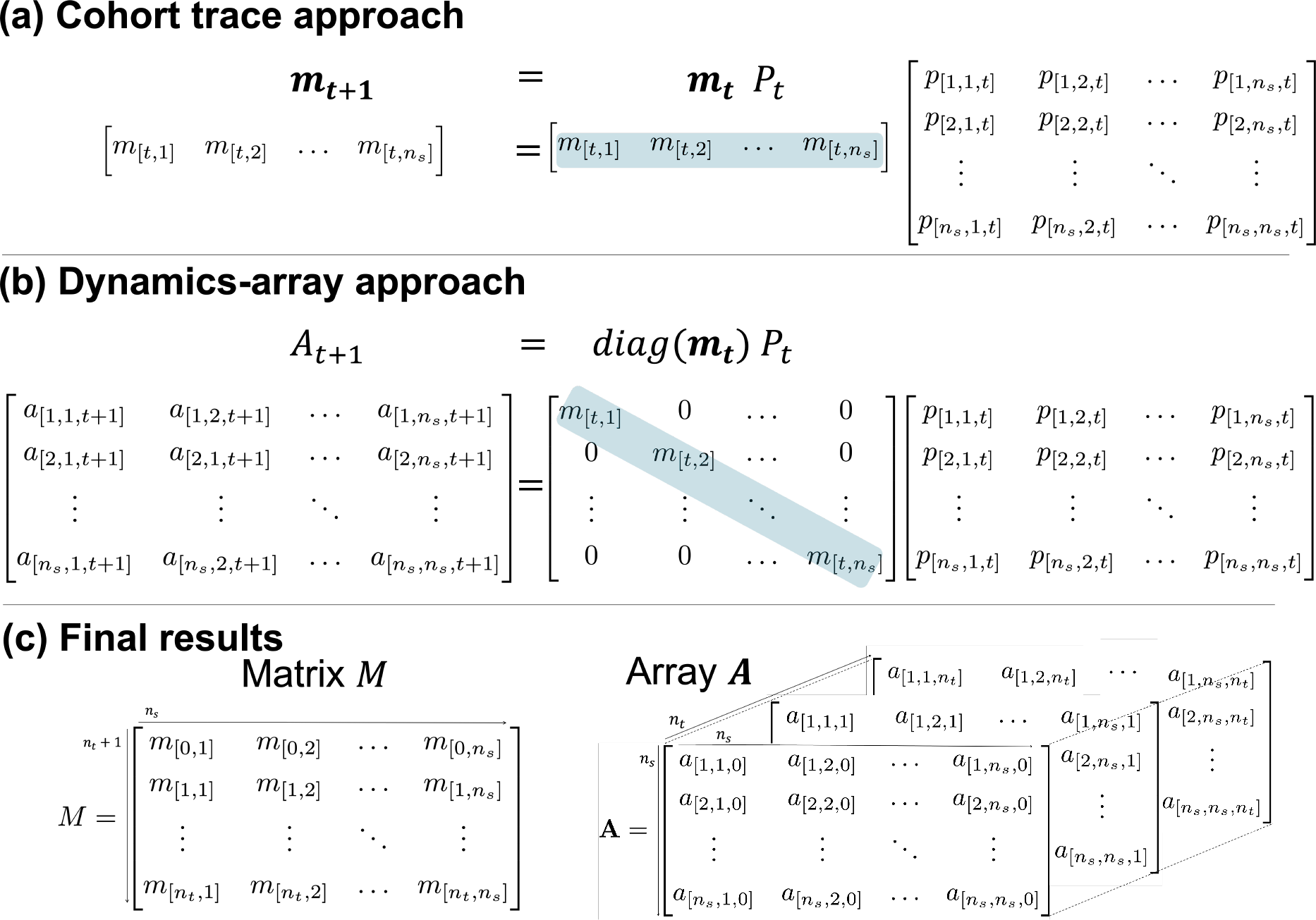
**(a)** The cohort trace approach computes row vector **m**_*t*+1_ of the cohort trace *M* that describes the distribution of the simulated cohort among the different health states at time *t* + 1. **m**_*t*+1_ results from multiplying the state vector **m**_*t*_ (gray) by the transition probability matrix *P*_*t*_. **(b)** The dynamics-array approach computes matrix *A*_*t* + 1_ containing information regarding the transition dynamics of the simulated cohort at time *t* + 1. The state vector **m**_*t*_ is highlighted (gray) to emphasize that the information in both approaches is identical. (**c**) Shows the resulting matrix *M* and array **A** of the approaches (a) and (b), respectively.

In R, it takes only a few lines of code to generate **A** complementary to *M* (Box 1).

**Box 1:**
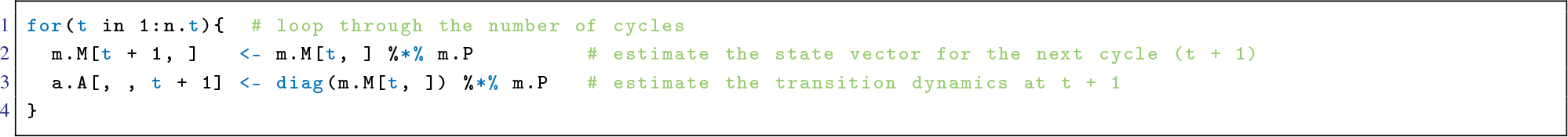
R code to iteratively generate the cohort trace *M* and the dynamics-array **A**.

The cohort trace *M* can be computed from **A** by obtaining the *t*-th row of *M*, **m**_*t*_, summing each of the columns of **A**_*t*_ as follows:

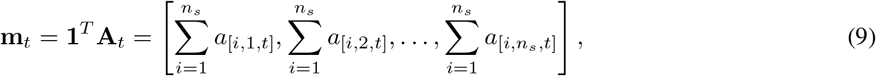

where **1** is a vector of ones of dimension *n*_*s*_ × 1. Although *M* can be obtained from **A** (avoiding computing *M* altogether), we prefer to compute both *M* and **A** simultaneously.

### Applying state and transition rewards

One of the main advantages of **A** over *M* is the ability to incorporate transition rewards. Here, we demonstrate how to apply both state and transition rewards (e.g. utilities or cost) to the cSTM by using the dynamics-array, **A**. Let *R*_*t*_ be a reward matrix of dimensions *n*_*s*_ × *n*_*s*_ that contains both state and transition rewards:

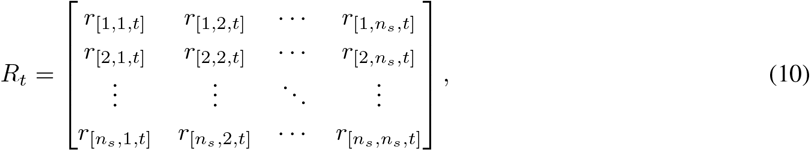

where *r*_[*i,j,t*]_ is the reward associated with transitioning from state *i* to state *j* at cycle *t*. When *j* = *i*, *r*_[*i,i,t*]_ is the reward associated with staying in the *i*-th health state at cycle *t*. That is, the off-diagonal entries of *R*_*t*_ store the transition rewards and the diagonal of *R*_*t*_ stores the state rewards for cycle *t*. The state and transition rewards can be applied to the model dynamics by element-wise multiplication between *A*_*t*_ and *R*_*t*_, indicated by the ⊙ sign, which produces the matrix of outputs at cycle *t*, *Y*_*t*_. Formally,

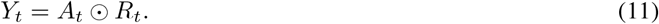

In R, applying these rewards required one additional line of code compared to Box 1, as shown in Box 2.

**Box 2:**
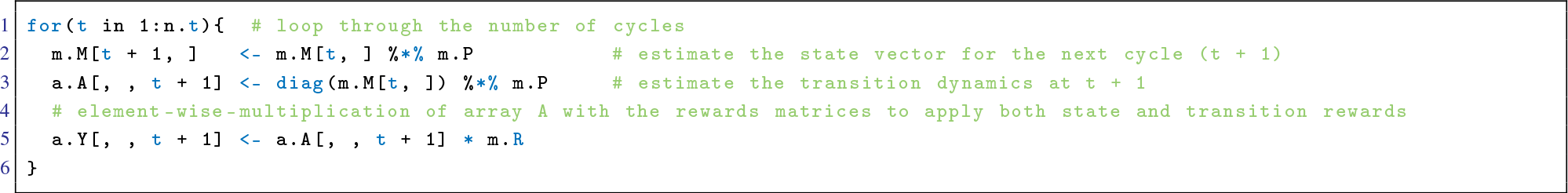
R code to apply time-invariant state and transition rewards to the model dynamics stored in array **A**.

The total rewards for each health state at cycle *t*, **r**_*t*_, is obtained by summing the rewards across all *j* = 1, …, *n*_*s*_ health states.

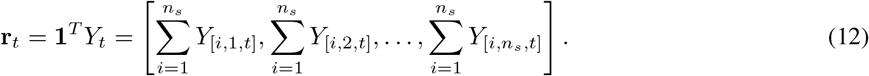

### Implementation in R using an illustrative example

To facilitate the implementation of the array approach, we demonstrate its use with a stylistic healthy-sick-dead 3-state time-homogeneous cSTM example coded in R [10]. The model is used to simulate a cohort of 70-year-old individuals to compute their expected costs and quality-adjusted life years (QALYs) accrued over their remaining lifetime accounting for several transition rewards. The explanation of the model and the R code can be found in the supplementary material and more detailed on GitHub (https://github.com/DARTH-git/state-transition-model-dynamics).

### Estimation of epidemiological measures

By obtaining **A**, it is possible to compute epidemiological outcomes that otherwise would not have been easily derived from *M*. For example, obtaining incidence and lifetime risk from *M* would require creating additional steps, variables or health states. Epidemiological outcomes could be used as outputs of simulation models for calibration or validation purposes. A full exposition of computing epidemiological measures from **A** is case-specific and is beyond the scope of this brief report. However, we illustrate the potential application of our approach by calculating a simple ratio from a generic cSTM.

Consider a cSTM with *n*_*s*_ > 3 health states. We are interested in calculating a ratio *e*_*t*_ of those that transition from health state 2 to health state *n*_*s*_ at cycle *t* to those that make this transition from health states 1, 2 and 3. Using the dynamics-array approach, the ratio *e*_*t*_ can be computed as

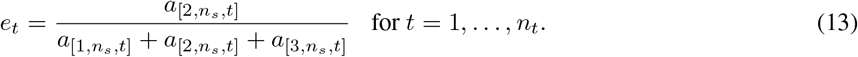

## Discussion

We propose a multidimensional-array approach to overcome a limitation of the cohort trace produced by cSTM in not being able to store transition dynamics. The practical application of our approach involves adding a simple step to the traditional cohort trace approach that stores all transitions among health states over time in multidimensional array **A**.

Traditionally, researchers have dealt with this limitation of the cSTM cohort trace by creating temporary health states that collect the state-to-state transition information. However, this solution can quickly complicate a model and result in an explosion of the number of health states. Using an individual-based microsimulation STM is another alternative [1], with considerable implications on computational time [11].

Another method that explicitly keeps track of state-to-state transitions is through a discretely integrated condition-event (DICE) simulation [8, 7]. DICE is a modeling technique that can free up some of the Markov restrictions that makes it possible to explicitly include many events occurring at various times. Although DICE simulation is a well-structured method, and the authors of the DICE papers provided very useful supplementary files to apply the method, we see the dynamics-array approach as a relatively simpler method to compute than DICE to overcome the limitation of the cohort trace on applying transition rewards and generating all the epidemiological outcomes of interest.

A potential limitation of the use of **A** is the additional computation needed when building the model. However, for many applications this may be a minor limitation given the matrix-based computational efficiency of current computers. Another potential limitation is the additional storage memory required to store **A**, which could become a limitation in systems with limited memory. This could be an issue for computationally complex models with multiple states. However, the benefit of using **A** for large models is that all the complexity in the model dynamics is summarized into a compact structure which makes it relatively simple to extract information or to apply new rewards without re-running the model.

In conclusion, structuring the output of cSTMs using the dynamics-array is an efficient, simple and convenient approach to summarize the model dynamics. This simple structure allows applying state and transition rewards and obtaining epidemiological measures, while still being able to obtain and display the conventional cohort trace.

## 1 Supplementary material: R code of the stylistic 3-state model

### Model description

We follow a cohort of healthy 70-year-old individuals over their remaining lifetime, using 30 annual cycles. The healthy individuals can transition to the sick health state, they can die or remain healthy. Sick individuals can fully recover, transitioning back to healthy, remain sick or die. Remaining in each of these health states is associated with some utilities and costs (the state rewards). In addition to these state rewards, transition dis-utilities and costs apply. Getting sick is associated with a sudden decrease of quality of life of 0.1. In addition, transitioning to dead incurs a one-time cost of $4,000. Both the state and transition rewards are constant over time. The R code to use the dynamics-array approach for this case example is shown below. All parameters of this model are fictitious, not based on a specific disease. On GitHub, we describe this simple 3-state example in more detail GitHub - (https://github.com/DARTH-git/state-transition-model-dynamics).

### R code

We recommend downloading this code from GitHub to avoid errors due to copying from the manuscript.

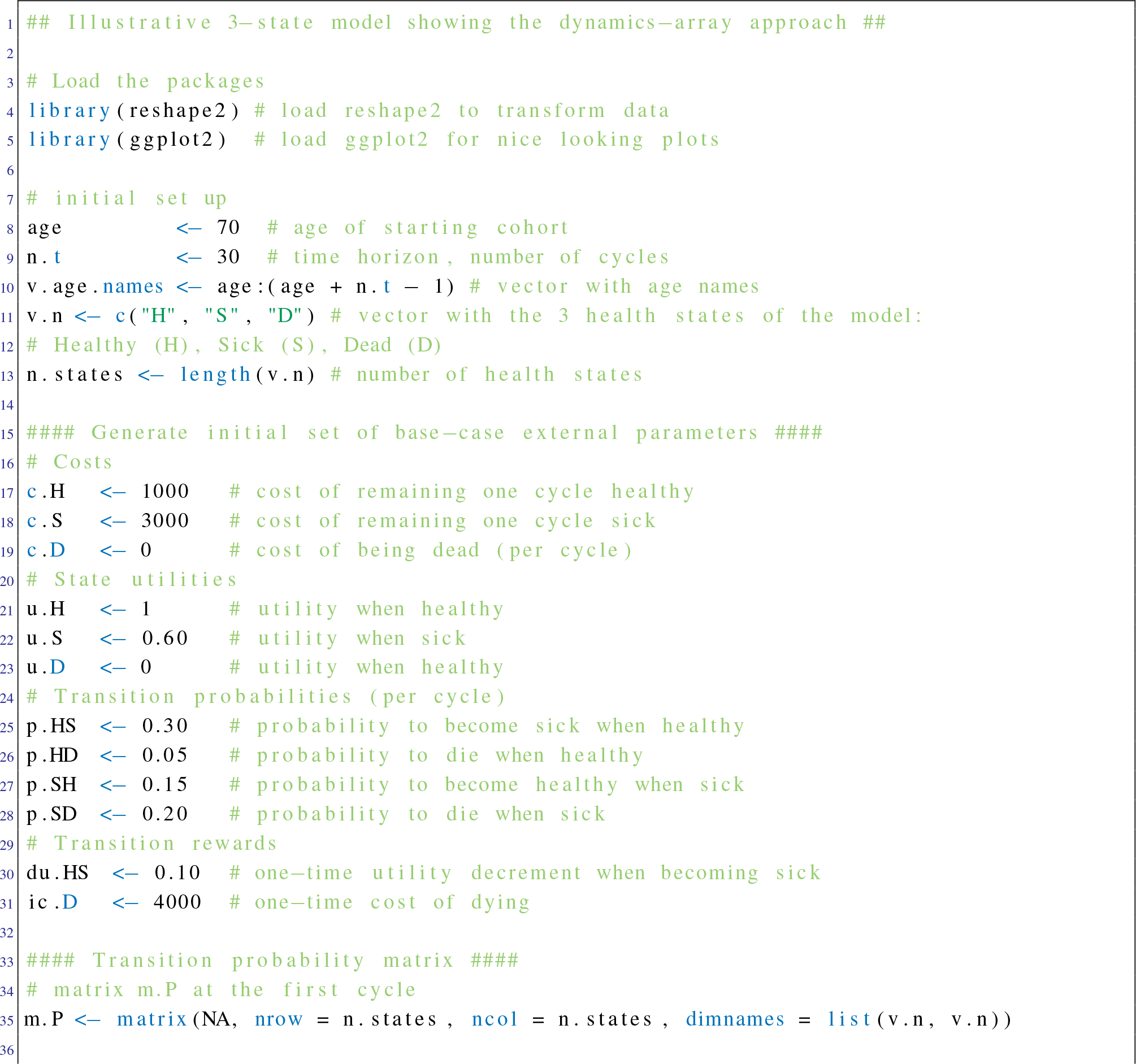
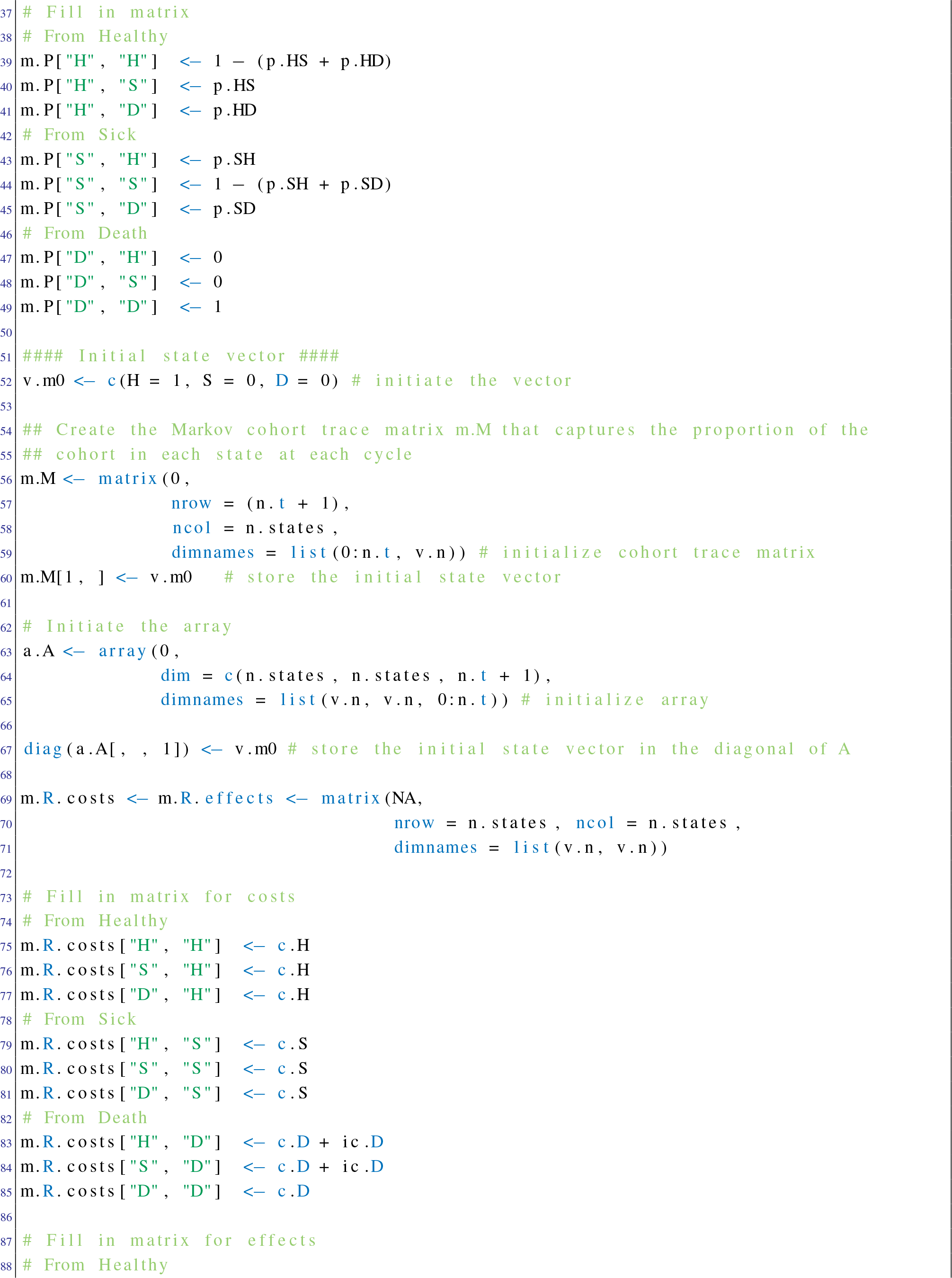
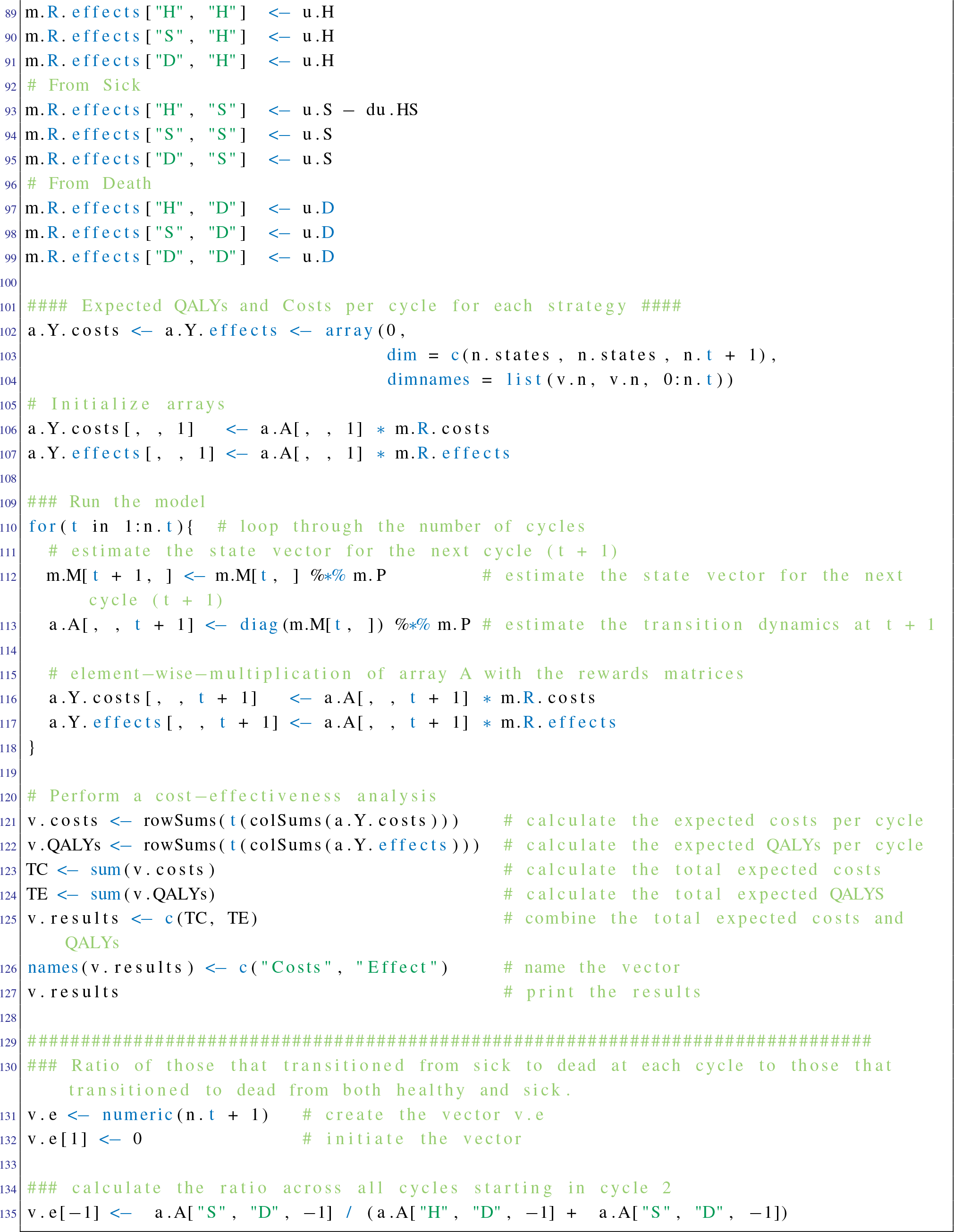

## References

[1] Siebert U, Alagoz O, Bayoumi AM, Jahn B, Owens DK, Cohen DJ and Kuntz KM. State-Transition Modeling: A Report of the ISPOR-SMDM Modeling Good Research Practices Task Force-3. Value Health, Sep-Oct;15(6): 812–20, 2012.

[2] Kuntz KM, Russell BL, Owens DK, Sanders GD, Trikalions TA and Salomon JA. Decision Models in Cost-Effectiveness Analysis. In Cost-Effectiveness in Health and Medicine, Neumann, PJ, Ganiats TG, Russell LB, Sanders GD, Siegel JE, Chapter 5, pages 105–36. Oxford University Press, New York, second edition, 2017.

[3] Beck JR and Pauker SG. The markov process in medical prognosis. Med Decis Making, Dec;3(4):419–58, 1983.

[4] Iskandar R. A theoretical foundation for state-transition cohort models in health decision analysis. PLoS One, Dec;13(12): 1–11, 2018.

[5] Sonnenberg FA and Beck JR. Markov models in medical decision making: A practical guide. Med Decis Making, Oct-Dec;13(4): 322–38, 1993.

[6] Bouter LM, Zielhuis GA and Zeegers MPA. Textbook of Epidemiology Bohn Stafleu van Loghum, 2018.

[7] Caro JJ and Möller J. Adding Events to a Markov Model Using DICE Simulation Med Decis Making, Feb;38(2): 235–45, 2018.

[8] Caro JJ Discretely Integrated Condition Event (DICE) Simulation for Pharmacoeconomics PharmacoEconomics, Jul;34(7): 665–72, 2016.

[9] Jalal HJ, Pechlivanoglou P, Krijkamp EM, Alarid-Escudero F, Enns EA and Hunink MGM. An Overview of R in Health Decision Sciences. Med Decis Making, Oct;37(7): 735–46, 2017.

[10] R Core Team. R: A Language and Environment for Statistical Computing. R Foundation for Statistical Computing, Vienna, Austria, 2013.

[11] Krijkamp EM, Alarid-Escudero F, Enns EA, Jalal HJ, Hunink MGM and Pechlivanoglou P. Microsimulation Modeling for Health Decision Sciences Using R: A Tutorial. Med Decis Making, Apr;38(3): 400–22, 2018.

